# Carbapenem heteroresistance of KPC-producing *Klebsiella pneumoniae* results from tolerance, persistence and resistance

**DOI:** 10.1101/2022.05.03.490393

**Authors:** Adriana Chiarelli, Nicolas Cabanel, Isabelle Rosinski-Chupin, Thomas Obadia, Raymond Ruimy, Thierry Naas, Rémy A. Bonnin, Philippe Glaser

## Abstract

Carbapenemase-producing *Klebsiella pneumoniae* (CPKp) have disseminated globally and represent a major threat in hospitals with few therapeutic options and high mortality rates. Isolates producing the carbapenemase KPC (KPC-Kp) might be classified as susceptible according to clinical breakpoints by antibiotic susceptibility testing (AST), allowing the use of imipenem or meropenem for treatment of infections. However, some KPC-Kp show heteroresistance (HR) to carbapenems, with colonies growing in the inhibition halo of agar-based AST. HR KPC-Kp have been associated with a higher risk of treatment failure. Here, we characterized the diversity of mechanisms behind HR to imipenem of these isolates. By analyzing a diverse collection of CPKp, we showed that HR is frequent among KPC-Kp. By monitoring single HR colony appearance using the ScanLag setup, we discriminated surviving cells in two subpopulations leading to a Gaussian-like distribution of early-appearing colonies, with a delayed emergence compared to colonies arising in the absence of antibiotics, and a long tail of late-appearing colonies. A subset of colonies showed a reduced growth rate. Characterization of surviving populations by AST and whole-genome sequencing of 333 colonies revealed a majority of parental genotypes and a broad landscape of genetic alterations in 28% of the colonies, including gene loss, DNA amplification and point mutations. This unveils the complexity of imipenem HR among KPC-Kp isolates, which involves tolerant and persistent cells, but also resistant bacteria. These observations contribute to a better understanding of reasons behind carbapenem treatment failure of KPC-Kp isolates.

**IMPORTANCE:** The ability of a bacterium to defeat antibiotics not only depends upon resistance, but also on tolerance and persistence, which allow a bacterial population to temporarily survive high drug doses. Carbapenems are antibiotics of last resort and *Klebsiella pneumoniae* isolates producing the carbapenemase KPC are a threat to hospitals, although they might remain susceptible to carbapenems. However, seemingly homogeneous populations of KPC-*K. pneumoniae* isolates frequently show varying degrees of susceptibility to carbapenem, i.e., a phenomenon called heteroresistance. We characterized bacteria surviving a high dose of imipenem, progressively degraded by the released carbapenemase, by monitoring the growth of the resulting colonies using the ScanLag system, their genome sequence and carbapenem susceptibility. We show that the observed phenotypic diversity combines tolerance, persistence and resistance making the treatment with high doses of carbapenems frequently inefficient.

## INTRODUCTION

Antibiotic resistance represents a global health concern. Several bacterial pathogens evolved multi-drug resistance (MDR), critically affecting their clinical management. Carbapenem-resistant Enterobacterales (CRE), including *Klebsiella pneumoniae*, were listed by the World Health Organization as critical priority pathogens for which there is an urgent need to develop new antibiotics (1). Carbapenem resistance may arise from the production of an extended-spectrum ß-lactamase (ESBL) or a cephalosporinase associated with decreased permeability due to mutations in porin genes and/or upregulation of efflux pumps, or resulting from the production of specific carbapenem-hydrolysing β-lactamases (carbapenemases) (2). Carbapenemase-producing *K. pneumoniae* (CPKp) is one of the most feared nosocomial pathogen worldwide (3). Carbapenemases belong to Ambler class A (KPC-type), class B (metallo-β-lactamases IMP, VIM- and NDM-types) or class D (OXA-48-like enzymes) of ß-lactamases (4) with variable occurrence according to geographical areas (5, 6). In clinical practice, CPKp may be treated with high doses of carbapenem depending on the minimum inhibitory concentration (MIC) (7). Indeed, production of a carbapenemase leads to a decreased susceptibility to carbapenems but can remain under clinical breakpoints. However, treatment effectiveness is extremely dependent on the accurate measurement of carbapenem MIC (8), and MIC determination is often challenged by variable resistance levels shown by CPKp, especially those producing the class A carbapenemase KPC (KPC-Kp) (9, 10).

A further challenge is represented by heteroresistance (HR) towards carbapenems. Antibiotic HR broadly describes heterogeneous susceptibility levels to an antibiotic within seemingly isogenic bacterial populations. This “bet-hedging” survival strategy confers long-term fitness benefit to populations in constantly fluctuating environments (11). In addition, as resistance is often a stepwise evolutionary process, such heterogeneity may precede genetic changes, and accelerate resistance development in the population (12, 13). Antibiotic HR itself corresponds to different situations according to the clonality of resistant subpopulations compared to the bulk of the population, the resistance level and the stability of the observed phenotype (14). Most of the studies addressing the HR issue adopt the Population Analysis Profiling (PAP) method, although following different parameters. Yet, there is a general consensus that a strain can be defined as HR when showing subpopulations growing at more than eight times the MIC of the main cell population (14).

Antibiotic HR has been largely reported among *K. pneumoniae* isolates towards colistin and corresponding to resistant subpopulation due to mutations modifying the LPS charge (15, 16). Although to a minor extent, HR has also been described towards carbapenems (17-20). Carbapenems HR was mostly observed among KPC-Kp (19-21). However, mechanisms mediating carbapenem HR in *K. pneumoniae* remain poorly characterized. Heteroresistant KPC-Kp strains survival upon exposure to high bactericidal concentrations of imipenem (IMP) was shown to result from the emergence of a resistant mutated subpopulations with decreased expression of the major outer membrane porin (OMP), OmpK36 (21, 22). Survival is likely favored in the first place by IMP degradation driven by KPC released into the culture medium, reminding of the phenomenon of indirect resistance, where an antibiotic resistant bacteria contribute to the growth of susceptible ones (23).

Here, we aimed at characterizing carbapenem HR in KPC-Kp clinical isolates and identifying metabolic and regulatory pathways underlying the diversity of susceptibility levels to high doses of carbapenems. We characterized phenotypically and genotypically a collection of 62 KPC-Kp isolates, in addition to 20 *K. pneumoniae* isolates producing other carbapenemases. A subset of KPC-Kp isolates was further selected to infer the dynamics of bacterial response to high doses of IMP at single colony level (24). Through the characterization of surviving subpopulations, we showed that HR to imipenem in KPC-Kp is a multifaceted phenomenon, which encompasses different mechanisms adopted by bacterial populations to survive drug exposure, namely tolerance, persistence and resistance. This approach also revealed a broad diversity of genetic alterations in diverse pathways associated with the observed phenotypes.

## RESULTS

### Diversity of carbapenem resistance and prevalence of heteroresistance among KPC-Kp

In order to characterize HR among KPC-Kp, we collected 62 KPC-Kp clinical isolates. For comparison, we included 20 clinical isolates expressing other carbapenemases and three clinical isolates without any carbapenemase as control (Table S1). Whole genome sequence (WGS) analysis revealed that the KPC-Kp isolates belong to 16 STs, with ST512 being the most prevalent (n = 19), followed by ST101 (n = 10) and ST258 (n = 8). Thirteen different capsular types were identified, with KL107 being the most represented one (n = 24) and mostly associated with ST512. In addition to the carbapenemase genes, the 82 CPKp isolates displayed diverse antibiotic resistance genes and porin mutations (Table S1). Defects in at least one of the major porin genes (*ompK35* and *ompK36*) were detected in 66% of isolates (n = 54/82) and, among them, 33% (n = 18/54) co-harbored an ESBL gene. In agreement with previous studies, CPKp ST512 isolates shared a truncated *ompK35* sequence and a G134D135 duplication in the sequence of the OmpK36 L3-loop, resulting in a narrower pore and yielding similar resistance levels as a porin inactivation (25). Likewise, CPKp ST258 isolates exhibited a truncated *ompK35* sequence and, two of them had the same *ompK36* GD duplication. The ten ST101 CPKp isolates shared a frameshift mutation in *ompK35* and nine of them also had a TD or GD duplication in *ompK36* and one a D duplication in OmpK36 L3-loop (Table S1).

The susceptibility levels to IMP and MEM were determined by Etest strip, as the broth microdilution method might be affected by carbapenem degradation (26). The carbapenemase-producing isolates exhibited a broad range of MICs, from 0.125 to >32 mg/L to MEM and 0.25 to >32 mg/L to IMP respectively (Table S1). All the IMP R isolates (n = 19) exhibited defects in the sequence of at least one major porin, with 95% (n=18/19) showing altered sequences of both *ompK35* and *ompK36*. All MEM R isolates (n = 38) but one showed at least one mutation in *ompK35*, with 89% having both *omp* genes disrupted (n = 34). A minority of MEM R isolates (9/38) and IMP R isolates (5/19) carried at least one ESBL gene. This observation suggested that higher carbapenem MICs did not result of the expression of an ESBL in the KPC-Kp isolates we have analyzed but mainly of reduced OM permeability.

We next evaluated HR of these isolates based on the appearance of colonies within the inhibition halo induced by the Etest antibiotic gradient strip (Fig. 1A). Isolates showing full contact with the Etest gradient strip (IMP, n = 3; MEM, n = 31) for which HR could not be detected were excluded from the analysis (Fig. 1A). According to the criteria defined in the Methods section, 73% (n = 43/59) and 51% (n = 17/31) of the KPC-Kp isolates were identified by Etest as HR or moderate HR to IMP and MEM, respectively (Fig. 1B, Table S1). A higher rate of IMP HR was detected for isolates otherwise classified as susceptible (78%) or intermediate (80%) based on their MIC compared to resistant ones (67%). In the case of MEM, HR was more frequent with isolates classified as intermediate (58%) (Fig. 1B). In addition, 71% of isolates carrying *bla*_KPC-2_ showed MEM HR isolates, in contrast with only 30% of those harboring *bla*_KPC-3_ (Fig. 1C). Using the same criteria, HR was also detected by Etest among NDM-producing *K. pneumoniae* (n = 3/5 HR to IMP 2/5 HR to MEM), but was not observed among the VIM- and OXA- and control isolates (Table S1).

**Fig. 1.**
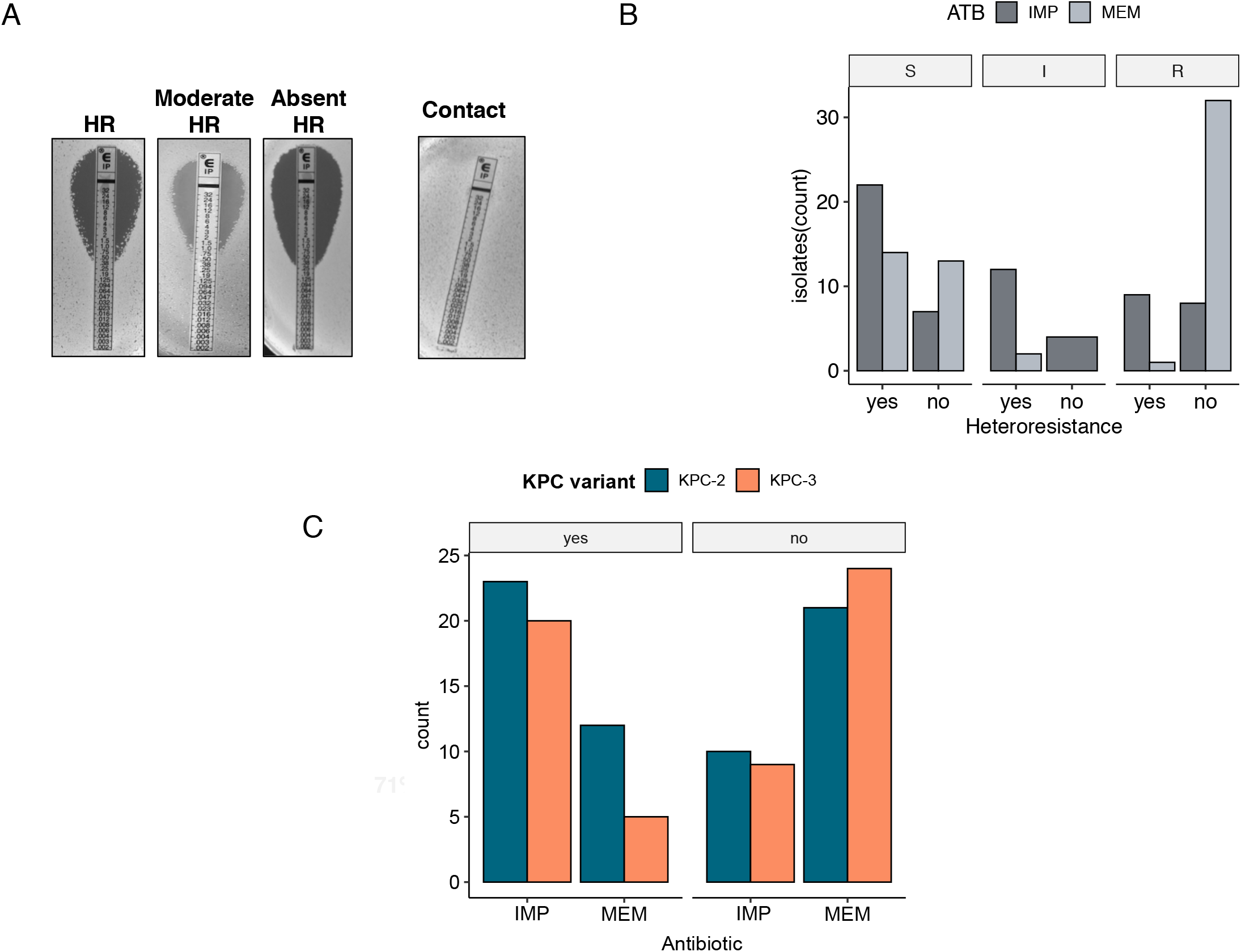
Diversity of resistance and heteroresistance. (A) Visualization of the HR profiles by Etest. Three different categories were distinguished (HR, moderate HR and absent HR). Isolates showing full contact with the antibiotic strip (on the right) were excluded from further analyses and classified as “HR not detectable” (Table S1); (**B)** Occurrence of HR among KPC isolates. SIR = Susceptible/Intermediate/Resistant. Bar colors: dark grey = imipenem (IMP), light grey = meropenem (MEM). Yes = presence of HR (including HR and moderate HR isolates), No = absence of HR. Isolates that showed full contact with the antibiotic strip (*n* = 31 for MEM and *n* = 3 for IMP) were not included; (**C)** Correlation between HR profile and type of KPC carbapenemase. Bar colors: Blue = KPC-2; Orange = KPC-3; yes for HR and no for non-HR.

Altogether, these results showed that our set of isolates harboring *bla*_KPC_ exhibited a broad range of resistance and HR levels.

### KPC-Kp isolates show subpopulations growing at high imipenem doses

Based on MIC values and HR data, we selected seven KPC-Kp in our set of isolates to characterize their response to IMP concentrations 4-, 8- and 10-fold their MICs, as determined by Etest. For comparison, two NDM-Kp, two OXA-Kp and two VIM-Kp were also analyzed (in bold in Table S1). To evaluate the impact of the growth phase on the survival of subpopulations to IMP, bacterial cultures in exponential (EP) and stationary phase (SP) were tested.

According to the definition of HR, isolates showing colonies at least at IMP 8xMIC were considered as heteroresistant. We detected HR, to varying degrees, in four KPC-Kp isolates: CNR146C9, CNR121E10, KP H15-21-6 and BIC-1 (Fig. 2A). The highest frequency of survival at 8- and 10-fold MIC of IMP was observed for CNR121E10, followed by CNR146C9, thus confirming their HR profile displayed by Etest. Interestingly, KP H15-21-6 was not HR by Etest, but showed surviving subpopulations by PAP. On the other hand, although NDM isolates showed a HR-like pattern by Etest, they did not show HR by PAP assay at 8-fold MIC of IMP. VIM- and OXA-isolates showed scarce or none ability to grow at 8-fold MIC IMP, in agreement with undetected HR profiles by Etest. Significant differences between EP and SP inoculum were observed for four out of the 13 isolates at 8-fold MIC (Fig. 2B). The two KPC-Kp (CNR146C9 and CNR121E10) and the VIM-Kp (CNR206A5) isolates showed a higher frequency of surviving subpopulations in SP, suggesting a persister-like subpopulation. However, the KP1 strain, producing VIM-1, displayed a higher number of surviving CFUs in EP than in SP. These observations suggested a strain-specific ability to cope differently with high doses of IMP, according to the physiological state of the bacterial population.

**Fig. 2.**
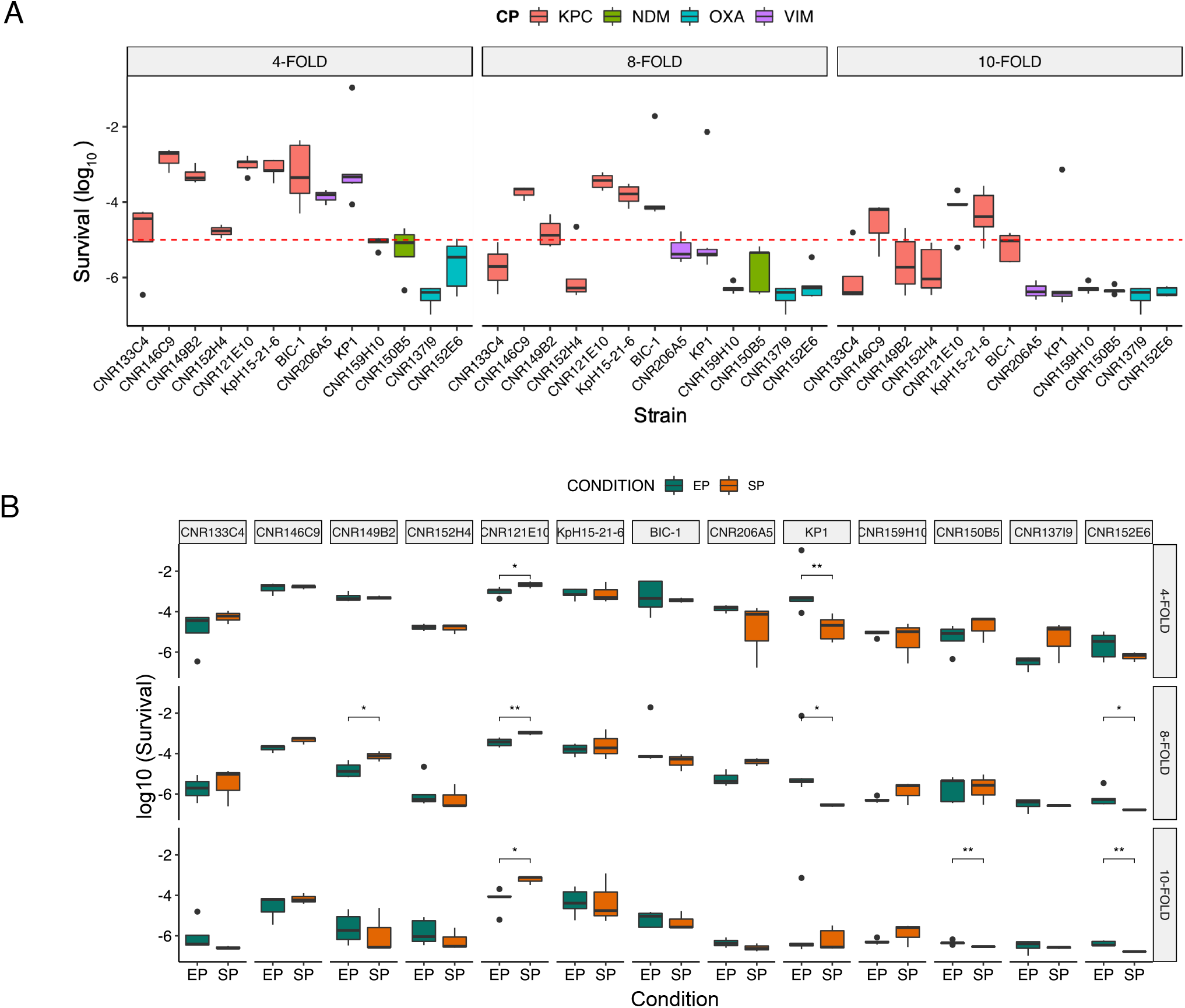
Population analysis profiling (PAP) of 13 CPKp clinical isolates. **(A)** Results were expressed in log_10_ of the frequency of survival at 4-FOLD, 8-FOLD, 10-FOLD the concentration of IMP relative to the MIC of the isolate as determined by Etest. Isolates names are indicated at the top of each chart. Red bars: KPC isolates; magenta bars: VIM isolates; green bars: NDM isolates; blue bars: OXA isolates. The red dashed line indicates the cutoff value (−5) above which counts were considered significant. (**B)** Impact of the bacterial growth phase on the survival to different doses of imipenem. Green bars indicate plating from exponentially growing cultures, whereas orange indicates plating from stationary phase bacterial cultures. Data were calculated from five independent experiments. Significant results were highlighted: * *p*-value ranging from 0.0159 to 0.0278; ** *p*-value ranging from 0.0025 to 0.093.

### Subpopulation growth dynamics

Differences in colony size were noticed on medium supplemented with 8-fold MIC IMP for CNR146C9, CNR121E10, Kp H15-21-6 and BIC-1 (Fig. S1A), indicating an heterogeneous growth of surviving subpopulations in the presence of IMP. To better characterize the dynamics by which subpopulations survive exposure to high IMP concentrations, we tracked appearance and growth of individual colonies in the presence of IMP by ScanLag (24). For comparison, we included in our analysis two KPC-Kp isolates showing scarce or none HR (CNR149B2 and CNR152H4). For these two isolates, colony behavior was investigated at 4xMIC IMP, as the number of CFUs at 8xMIC was below our detection threshold (less or equal than 10^2^ CFU/ml). Imipenem MICs for the six selected KPC-Kp isolates ranged from 0.25 to 1.5 mg/L (Table S1). By using ScanLag, we quantified the distribution of colony time of appearance (ToA) of the six isolates over 48 hours onto drug-free and IMP-containing media (Fig. 3A). In the absence of IMP, we observed a narrow normal distribution of ToA for all the isolates, with mean values ranging from 426 to 518 minutes (Table S2, Fig. 3A; cyan blue bars). By contrast, the distribution of colony ToA onto media supplemented with IMP was heterogeneous with a main peak and a tail of varying length, particularly visible in CNR146C9, CNR152H4 and CNR121E10 (Fig. 3A, red bars).

**Fig 3.**
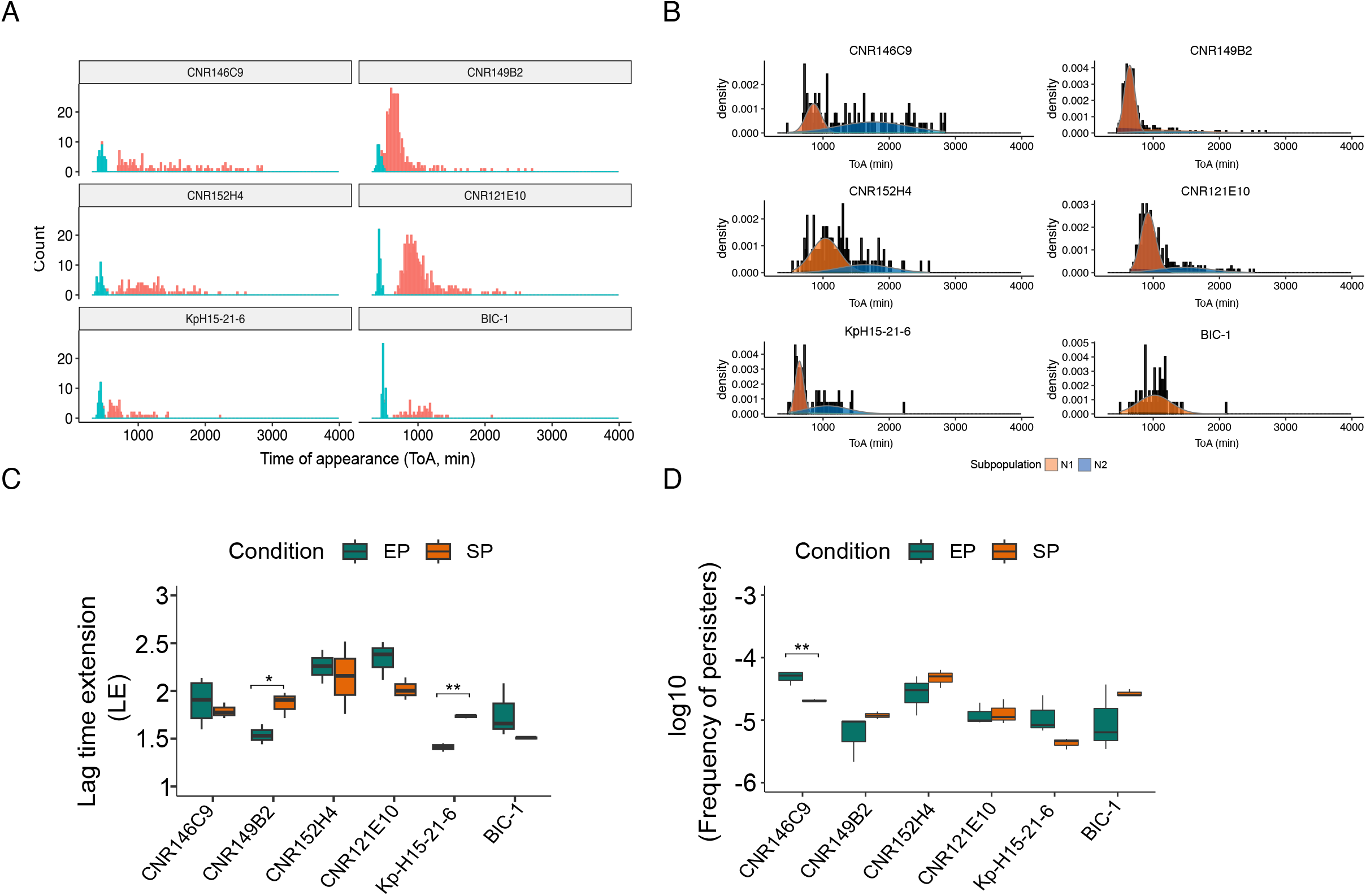
Heterogeneous survival of KPC-Kp to high concentrations of IMP. **(A)** Histograms of single-colony time of appearance from exponentially growing cultures. Colony appearance time was monitored every 20 minute over 48-hours incubation on plate. ToA = Time of Appearance; cyan blue bars represent colonies appearing on drug-free media, whereas red bars indicate colonies growing onto IMP-containing agar plates. The data are an overlap of at least three independent replicates per each isolate and each condition. (**B)** Fit of two-mixture component model to the distributions of ToA for each isolates after plating from exponential phase cultures. In red, population 1 (N1), and in blue the population 2 (N2). The representation is an overlap of at least three independent replicates per isolate and per condition. ToA data are expressed as density rather than count. Estimated parameters of each subpopulations and replicates are given in Table S2. Note that, for the isolate BIC-1, the model failed to fit a normal distribution to the tail. (**C)** Comparison of Lag Time Extension (LE) of N1 subpopulations from the six isolates plated from EP and SP cultures. Only significant differences were highlighted (* corresponds to *p*-value = 0.033, ** corresponds to *p*-value = 0.0025). LE was calculated as the mean ToA value of N1 population over the mean ToA value of untreated population. **(D)** Frequency of persisters calculated according to the methodology proposed by Balaban et al (28) from at least three replicates (data are available in Table S2). Colonies appearing after the cutoff value (calculated as ToA > μ+3Σ, where μ= estimated ToA mean for N1 and Σ = estimated standard deviations) were defined as persisters. Frequencies were calculated as the proportion of CFU/ml of each subpopulation over the total number of CFU/ml of the initial inoculum. The frequency of persisters was significantly higher in exponential phase rather than in stationary phase for CNR146C9 isolate (**, *p*-value = 0.0038).

The bimodal distribution suggested for the six strains the existence of two subpopulations: an early appearing N1 population corresponding to the main peak of ToA, and a late-appearing N2 population indicated by the tail. To characterize the observed heterogeneity, we attempted to fit the data to a two-component mixed model using ToA as index. This approach allowed us to quantify the proportions of the N1 and N2 subpopulations across the six isolates and the heterogeneous delay in growth (here referred to as lag time extension, LE) of the N1 population relative to the colonies appearing on drug-free media (Fig. 3B, Table S2). To estimate the influence of the bacterial growth phase on the survival rate and on the frequency of subpopulations N1 and N2, we also tracked the colony ToA upon spreading stationary phase (SP) cultures of the six isolates on the drug-free and IMP-supplemented agar media.

N1 subpopulation from isolates CNR149B2, Kp H15-21-6 and BIC-1 grown up to EP showed an earlier appearance (LE <2) in comparison with the other three isolates, with LE ≥ 2 (Fig. 3C, green boxplots). Since SP cultures have been shown to be more tolerant to ß-lactams (27), we expected to observe an extended delay in growth for SP cultures exposed to IMP. However, by quantifying the LE of N1 subpopulations, we could detect a significantly longer delay only for CNR149B2 and Kp H15-21-6 in SP compared to EP (Fig. 3C).

As previously proposed (28), colonies that appeared later than three standard deviations away from the mean of the main peak distribution (subpopulation N1) were hypothesized as derivative of persister cells, restoring growth only upon gradual exhaustion of IMP below the MIC. We estimated their frequency for the six isolates after plating on 8-fold MIC IMP (4-fold for strains CNR133C4 and CNR152H4) from exponential or stationary phase cultures. The highest frequency of persisters in exponential phase was observed for CNR146C9, whereas BIC-1 exhibited the lowest frequency (Fig. 3D). We did not detect any significant difference in the proportions of persisters between the EP and SP, except for CNR146C9 (p-value = 0.0038), which surprisingly showed a lower occurrence of persisters in SP, in contrast with the increased survival of SP grown bacteria as observed by PAP (Fig. 3D, Table S2).

Overall, we observed heterogeneous growth resumption among the selected isolates when plated on high concentrations of IMP (4-fold MIC or 8-fold MIC), which suggested the co-existence of at least two distinct phenotypes in the surviving populations. Both the growth delay (measured as lag time extension) and the proportion of the subpopulations N1 and N2 were strain-specific, likely linked to differences in the genomic backgrounds of the isolates.

### Genetic heterogeneity associated with the observed heteroresistance

Antibiotics HR refers to a phenotypically heterogeneous subpopulation within a seemingly isogenic population. However, increased survival might be also linked to genetic alterations. We selected CNR146C9 for in-depth genetic analysis of the HR populations and to better discriminate the events responsible for enhanced survival. To obtain a high quality reference genome, we determined its whole genome sequence by combining long- and short-read sequencing. CNR146C9 hosts two plasmids carrying *bla*_CTX-M-15_ and *bla*_KPC-3_ of 237,007 bp and 110,489 bp, respectively and a circular 52,608 bp-long extrachromosomal phage, previously reported in *K. pneumoniae* ST307 isolates (29) (Table S3).

Based on the ToA and colony growth rates as visualized by ScanLag (Fig. 3A), we classified surviving colonies according to three patterns: A, B and C (Fig. S1B). Patterns A and B emerged before the cut-off for persisters (on average 1382 min) and globally correspond to the subpopulation N1. Pattern A included colonies showing wild-type or near wild-type growth rates, whereas pattern B corresponded to colonies with reduced growth rates, as observed by ScanLag. Pattern C colonies emerged after the cut-off for persisters, showing a wild-type growth rate but delayed and variable ToA. They corresponded to the N2 subpopulation, the tail of the ToA distribution. A total of 333 colonies (167 of pattern A, 30 of pattern B and 136 of pattern C) were submitted right after isolation and 2 hour-growth in LB broth to WGS and IMP and MEM MIC determination by Etest. Ninety-four colonies showed at least one genetic modification, including single-nucleotide variations (SNVs), insertions, deletions of bases, plasmid/phage loss and amplification (Fig. 4A) affecting a broad range of functions (Fig. 4B).

**Fig. 4.**
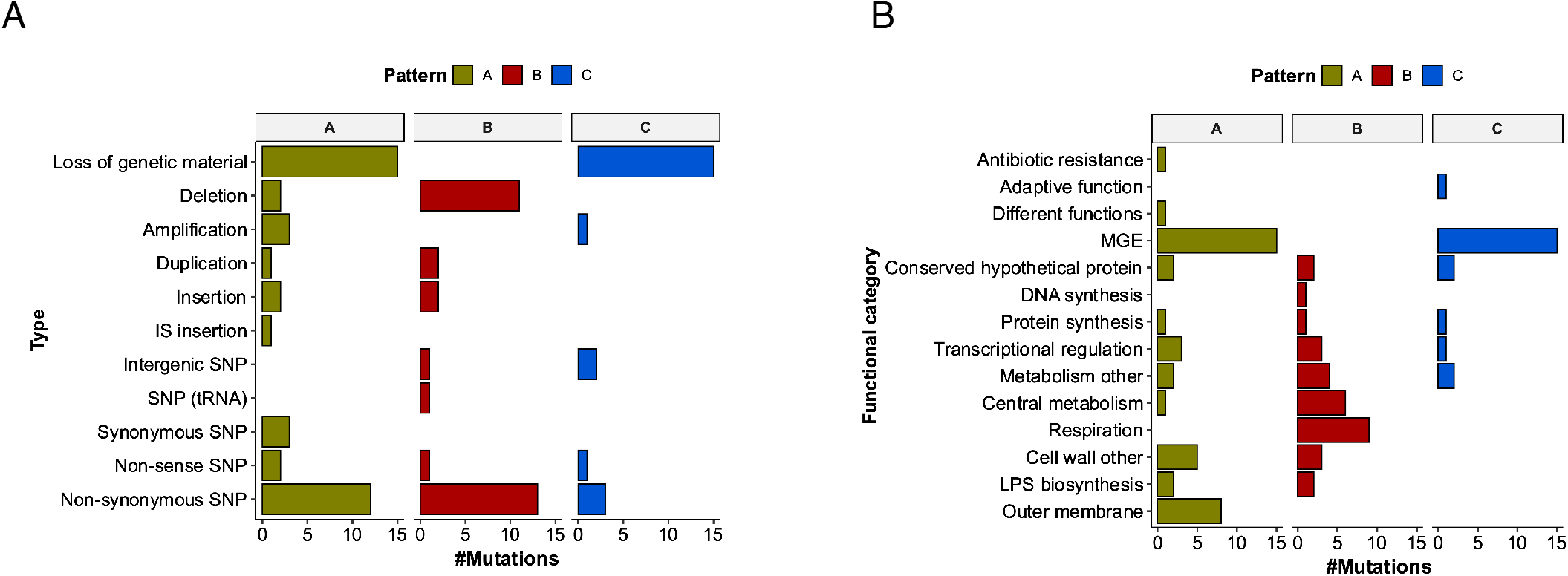
Diversity of genetic events detected among colonies from patterns A, B and C. **(A)** Diversity of genetic events as identified by using Breseq. **(B)** Functional categories of loci affected by the genetic events as identified in Tables S3, S4 and S5. The three classes of colonies as defined by their ToA and growth rate, A, B and C are indicated in green, red and blue respectively.

### Pattern A colonies result from tolerance or resistance to IMP

For most of the A-type colonies (128/167), we did not detect any genetic change. These colonies exhibited MIC values for MEM and IMP and an HR profile determined by Etest similar to the parental strain CNR146C9, suggesting they have survived in the presence of IMP until it reaches a concentration lower than the MIC. Thirty-nine A-isolates showed genetic alterations (Table S4). Most of them were not associated with a decreased susceptibility to both carbapenems (63%). The most recurrent event was the loss of the 52,608 bp extrachromosomal phage (n=12). One ICE and part of the *tra* region from the CTX-M-15 plasmid (n=3) were also lost. Point mutations were also detected among those colonies and affecting different functions like a nonsense mutation in *lysS* encoding one of the two tRNA synthase gene or a mutation in *rpsC* encoding the ribosomal protein S3. Some of these diverse genetic events might have been selected and contributed to longer IMP survival. On the other hand, 15 colonies showed a decreased susceptibility to IMP and MEM ranging from 1.5 mg/L to >32 mg/L. Strikingly, all mutations detected in these colonies except one were associated with alterations of the cell envelope. Mutations leading to the highest MICs affected the OM permeability, with three clones mutated in *ompK36*. However, non-canonical mutations affecting the Bam pathway of insertion of OMP into the outer membrane OM (30) were also selected. Three clones showed a truncated *bamB* gene, one was mutated in *bamA* (N448Y), which is essential, and one isolate showed an IS*Kpn25* insertion six bp upstream of the start codon of the *surA* gene encoding the major chaperon molecule contributing to the translocation of OMP across the periplasm (31). In addition, three clones show mutations affecting LPS biosynthesis and three others mutations in regulatory systems affecting cell wall components. Among these last isolates, AC90-A carried two mutations in the *cpxAR* locus encoding a two component regulatory system sensing OM stresses (32, 33): a four codons deletion (pos. 20 to 23) in the sensor *cpxA*, present in 100% of the reads and a R205W mutation in *cpxR* present in 95% of the reads. This clone is heterogeneous with dominant large colonies, mutated in *cpxR* and *cpxA* showing WT MICs to IMP and MEM and few highly resistant small colonies, only mutated in *cpxA* (Table S4). The only mutation associated with reduced susceptibility but not affecting the cell envelope occurred in the *lipA* gene, involved in lipoyl cofactor biosynthesis. Lipoyl is a cofactor of pyruvate and 2-ketoglutarate dehydrogenases, key enzymes of glycolysis and the tricarboxylic acid (TCA) cycle. This metabolic mutation reminds the type of mutations from the B pattern (see below).

Heteroresistance has been shown to result frequently from gene amplification leading to an increased gene expression (34). We systematically searched for DNA amplifications in the 333 genomes. We identified three cases of genome amplification among the A-type isolates (Table S4). In strain AC376, we observed a 89,584 bp ectopic duplication encompassing the *dcw* locus encoding major functions for cell wall synthesis and cell division, the additional copy being inserted upstream the transcriptional regulatory gene *qseB* (Fig. 5A). Interestingly, this isolate showed a 16-fold decreased IMP susceptibility but almost no effect on MEM susceptibility.

**Fig 5.**
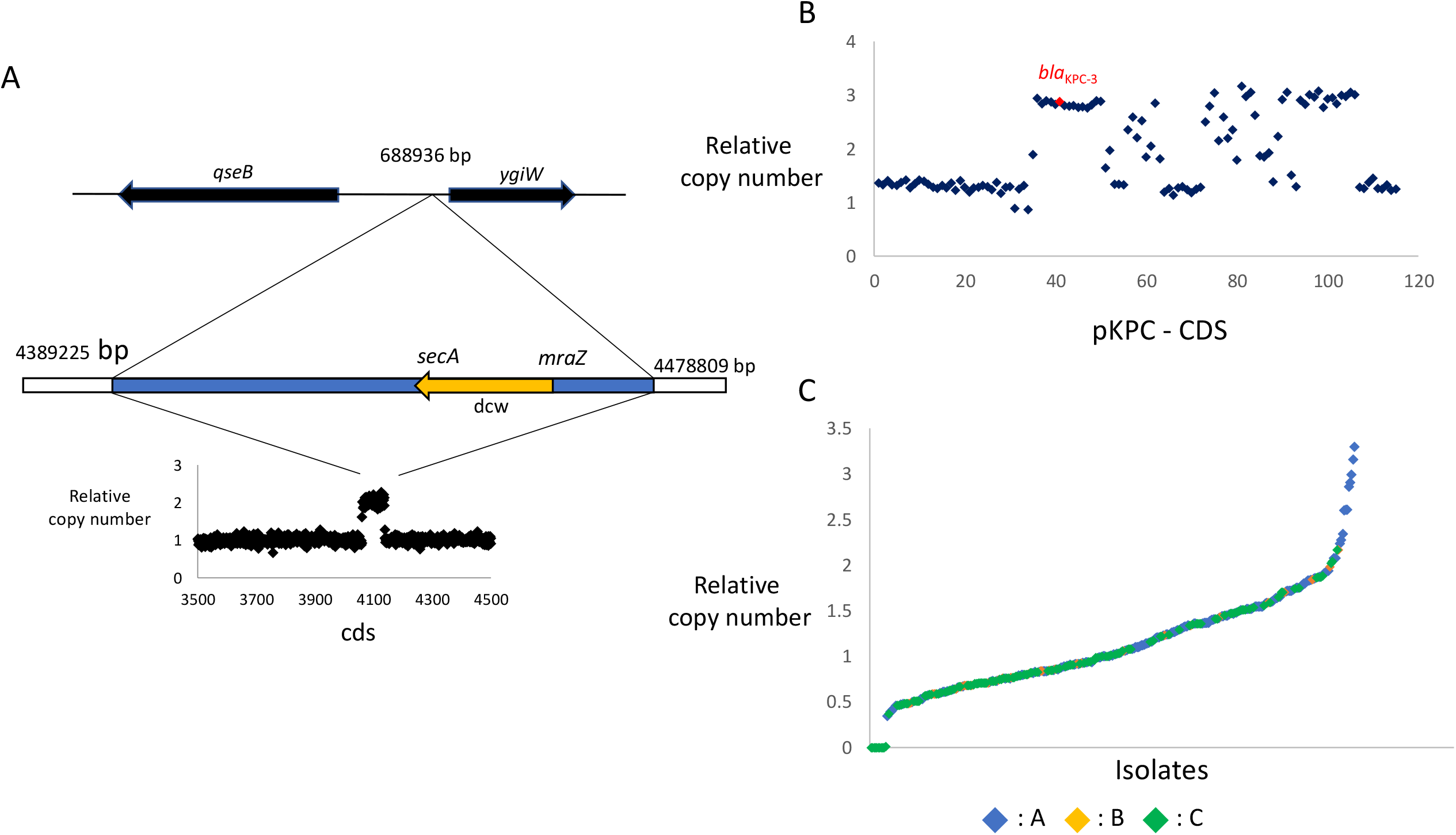
DNA amplification and pKPC copy numbers in pattern A isolates. (**A)** Duplication of a 89584 bp region in strain AC376-A. The relative copy number was estimated compared to the parental stain CNR146C9 based on the count of number of reads counted per gene. The duplicated region is in blue and contains as candidate locus contributing to increased IMP MIC the division and cell wall (dcw) locus (in orange). The insertion of the duplicated copy occurred 11 bp upstream *ygiW* with a 5 bp (GAGAA) duplication. *ygiW* expression is activated by the two component regulatory system QseBC encoded by two genes in the opposite orientation. Limits of the duplicated region and insertion sites were deduced from chimeric reads as identified by using Breseq (62). **(B)** Duplication of a c.a. 68 kb region within the pKPC plasmid in strain AC150-A. Average copy number of the 115 CDS of pKPC was calculated relative to the corresponding average value for all isolates except those having lost pKPC. The central part of the duplicated region is highly similar to a pCTX-M-15 DNA regions lost in this strain and explaining the variable and lower copy number as quantified by counting mapping reads (C) Average copy number of pKPC CDSs compared to the average copy number of pKPC CDS across all isolates, except those having lost pKPC. In blue, A isolates; in orange, B isolates and in green, C isolates.

Strain AC150-A showed a c.a. 68 kb-long duplication within the pKPC plasmid and encompassing *bla*_KPC-3_ (Fig. 5B). This duplication might be responsible for a doubling of the MICs for IMP and MEM of this strain. On the other hand, the gene coverage of pKPC was variable across all isolates with an average normalized coverage ranging from 0.39 to 2.62 excluding isolates which have lost pKPC. The 11 isolates with the highest pKPC coverage (ranging from 1.88 to 2.62-folds higher than the average) all belong to the A class. However, except AC150-A, none showed an increased MIC to IMP and MEM as quantified by E-test.

### Colonies of B-pattern are mainly small colony variants

Twenty-two out of the 30 B-pattern colonies here analyzed showed an increased MIC for both carbapenems. The eight remaining isolates might have been misclassified as pattern B or the phenotype was highly unstable, as after re-isolation they did not phenotypically differentiate from the parental strains (Table S5). We identified mutations in 28/30 isolates (Table S5) and 67% occurred in genes with metabolic functions. The most recurrent mutations affected respiration with mutations found in the coenzyme Q synthesis genes *ubiA* (n=2), *ubiB* (n=2), *ubiD* and *ispA* (n=3), in the heme biosynthesic gene *hemA* and in genes involved in the TCA cycle (a deletion encompassing *lpdA* and *aceF*; *icd*, encoding isocitrate dehydrogenase (n=2) and *acnB*, encoding the aconitate hydratase (n=2)). One isolate showed inactivation of the gene *ptsI* from the central phosphotransferase system (PTS), contributing to carbon source import and catabolic regulation in *E. coli* (35). In the presence of oxygen, these mutants showed tiny colonies with fastidious growth, which are features of small colony variant (SCV) (36). Under these conditions, they also showed an increased resistance to IMP and MEM. However, under anaerobiosis, these phenotypic traits were reversed for respiratory and TCA mutants. Therefore the metabolic defect was likely responsible for the phenotype (Table S5). Two mutations affected lipid metabolism: *pgpC* encoding a phosphatidylglycerophosphatase C and *hldE*, encoding a bifunctional kinase/adenylyltransferase involved in the inner core LPS biosynthesis (37). One isolate mutated in *cysB* showed an increased MICs only towards IMP. Interestingly, *cysB* mutation is the major cause of mecillinam resistance in clinical isolates of *E. coli* (38). One isolate (AC171-B) mutated in the tRNA^Ser^(gct) encoding gene showed a moderate decrease in susceptibility to both carbapenems and a strong growth defect, possibly due to altered protein synthesis. Globally, pattern B corresponded to resistant bacteria with mutations in *K. pneumoniae* core resistome and associated with decreased growth rate, as observed by ScanLag.

### Pattern C colonies likely correspond to persisters

Unlike colonies from pattern A and B, none of the pattern C colonies showed a significant increase in the MICs to carbapenems and 86% (n=117/136) did not show any detectable genetic modification. Unexpectedly, ten colonies showed an increased susceptibility to the two carbapenems due to the loss of the pKPC plasmid (Table S6). As observed for pattern A colonies, four patterns C colonies showed loss of the phage, whereas one clone (AC158-C) lost an integrated prophage. In one isolate (AC362-C), we observed a 3-fold copy number increase of a 51 kb long region from the pCTX-M plasmid bracketed by two IS*5075* (Fig. S2). This genomic island contains numerous adaptive functions for glycogen biosynthesis, lactose catabolism, the Fe(3+) dicitrate transport system (*fec* locus) and a glutathione transport system (29). Increased expression of these genes might have contributed to the enhanced survival as persistent cell. Two colonies showed intergenic mutations, while one nonsense mutation inactivates the type VI secretion system protein TssG. Whether these mutations contribute to the persistent phenotype remain to be determined.

The late appearance of the colonies and the nature of the seldom genetic events observed that are not contributing to ß-lactam resistance are in agreement with colonies resulting from persistent cells. These bacteria escaped the imipenem bactericidal effect until its concentration became lower than CNRR146C9 IMP MIC (0.5 mg/L) or until it is even below the MIC of a strain devoid of *bla*_KPC-3_ (0.19 mg/L).

## DISCUSSION

Although the expression of the KPC carbapenemase confers reduced susceptibility to carbapenems, MICs for imipenem are frequently below the breakpoint for susceptibility (1 mg/L, EUCAST), as we have observed in the set of 62 KPC-Kp clinical isolates analyzed in this work. Such MICs are theoretically compatible with the usage of IMP to treat those isolates, but with an increased risk of resistance selection and treatment escape (7). Moreover, occurrence of HR to IMP may further complicate the management of KPC-Kp isolates in clinical practice.

In addition to variable MICs, the KPC isolates here analyzed showed different levels of HR towards IMP and MEM. The proportion of isolates showing HR to IMP was higher than those with an HR profile to MEM. These two carbapenems share similar mode of action by targeting mainly PBP2, but also binding to PBP1a PBP1b. In addition, MEM shows a higher affinity to PBP3 (39), explaining differences in the morphological impact of those two carbapenems on *E. coli* cells (40). On the other hand, IMP is a smaller molecule than MEM and shows a rapid influx through the OM (41). More importantly, under laboratory conditions, IMP is less stable than MEM and is spontaneously degraded (41, 42), likely contributing to the higher rate of HR towards IMP comparted to MEM. We observed a higher proportion of HR towards MEM of isolates expressing KPC-2 compared to those expressing KPC-3 (Fig. 1C), which might be linked to a higher stability (longer half-life) of KPC-2 (43) contributing to a extended degradation of MEM in the solid medium.

HR has been defined as the occurrence of a bacterial subpopulation surviving treatment at 8x the MIC of the drug (14). In our PAP analyses, at this antibiotic concentration, frequency of survival ranged from 5.3 × 10^−6^ to 4.76 × 10^−4^. This low frequency challenges the harvest of the surviving population, limiting the use of microscopy for single cell analysis on agarose-pad or microfluidics device such as the mother machine (44). Previous studies addressing HR in KPC-Kp were performed in bulk, following addition of IMP at concentrations higher than the MIC leading to the selection of resistant mutants in the porin gene *ompK36* (21, 22). Here, we applied the ScanLag system (24) to analyze the individual fate of surviving bacteria forming colonies on media supplemented with IMP at 8xMIC or 4xMIC. For the six KPC-Kp isolates analyzed, ScanLag revealed a bimodal profile of colony appearance in the presence of IMP: a Gaussian peak, delayed compared to the peak of appearance of the colonies not exposed to IMP, followed by a tail of late appearing colonies. In addition to the delayed time of appearance, some colonies showed a slower growth rate. Compared to other antibiotics, a specific parameter of HR to ß-lactams in ß-lactamase producing bacteria is the degradation of the antibiotic in the medium. The first Gaussian peak of the surviving populations N1 is consistent with what has been defined as tolerant by adjusting lag time (45). Similarly, the late appearing colonies (N2) in the tail of the distribution, emerging under conditions where the antibiotic is gradually degraded would result from persister cells (28, 45). In agreement with these hypotheses, the vast majority of colonies from N1 and N2 populations of strain CNR146C9 reproduced the same susceptibility and HR phenotype as the parental strain. Comparisons of the six clinical isolates showed different profiles and effects of plating EP or SP bacteria (Fig. 3B and 3D), reflecting different levels of tolerance and persistence towards IMP. Such differences in drug survival are likely due to the genomic context, which remains to be fully investigated. As characterized in experimental models of evolution (46, 47), these features might have been selected in synergy with the selection of antibiotic resistance following drug treatments in the host.

One limitation of ScanLag is the absence of information on the first generations before the appearance of a visible colony. The observed delay might be related to an extended lag phase or to a transient slow growth phase reducing susceptibility. We speculated that combining genomic and phenotypic analyses of surviving colonies at 8xMIC would provide additional information on surviving populations. Most of the clones from profile A and C (from N1 and N2 populations respectively) showed a phenotype similar to the parental strain and did not exhibit any genetic events, which is in agreement with tolerance and persistence mechanisms, respectively. The frequent loss of pKPC plasmid in ten of the C-isolates was intriguing. Indeed, no plasmid loss was detected among 196 colonies of the parental strain CNR146C9 growing on antibiotic-free medium. The pKPC carries functions contributing to its stability including a *vapBC* family toxin/antitoxin (TA) locus. VapC toxins inhibit translation by an RNAse activity (48). In *E. coli*, the VapBC TA system was previously reported to contribute to increased rate of persistence (49) and a *vapB* mutant was associated with an extension of lag-time and higher drug tolerance in *E. coli* exposed to intermittent high doses of ampicillin (45). We hypothesized that the loss of pKPC might lead to a transient activation of VapC resulting of the instability of VapB and a growth arrest contributing to the persistent state. Therefore, pKPC loss would be under positive selection in the set-up we have used, supporting the hypothesis that C-type colonies arose mainly from persisters. Interestingly, loss of the KPC-plasmid was previously observed in clinical KPC-Kp isolates (50). In this context, the host response or antibiotic treatments might have contributed to a similar selection of persister cells. However, we cannot rule out the possibility that antibiotic stress contributed to plasmid loss. Among the diverse mutations identified not affecting IMP MIC, some might have occurred by chance. However, they might also have been selected as contributing to persistence or to an increased tolerance like mutation in the lysine tRNA synthetase *lysS*.

In addition to mutants with similar MICs as the parental strains, we observed an extremely broad range of mutations leading to increased MIC towards IMP, and, in most cases, also to MEM. Resistant colonies could be classified according to their fitness based on their growth rate, as determined by ScanLag (Fig. S1B). Most isolates showing the less affected fitness under the growth condition used (pattern A) were affected in OM permeability, with mutations in the porin gene *ompK36*, but also in the Bam system and in the major chaperon involved in the insertion of OMP in the OM. We also observed collateral effect of such mutations, like a hypermucoid phenotype for the *bamA* mutant or increased mutation rate in the *polB* isolate. Some mutations involving for instance regulatory systems and disrupting LPS biosynthesis likely affected the OM permeability indirectly (51). On the other hand, mutations from the B pattern showed a strong fitness cost and are reminiscent of SCV. SCV have been described in a broad range of species and are frequently associated with a decreased antibiotic susceptibility (36). However, there are only few reports of SCV in *K. pneumoniae*: in a bone and joint infection (52) and associated with colistin HR in biofilm (53). In both cases, the genetic bases for the SCV phenotype were not identified. As reported in other species, most mutations we have characterized affected aerobic respiration (quinone and heme biosynthesis) and the central metabolism (TCA cycle and glycolysis). Interestingly, both the slow growth and the increased MICs were fully or partially reversed in anaerobic conditions (Table S5). The combination of high resistance in the presence of oxygen and normal growth in its absence might contribute to the selection of such mutation during treatment. However, due to their slow growth under laboratory conditions, they might have been overlooked during diagnosis.

The increased MICs observed among HR colonies was shown to be frequently unstable and lost following passages in antibiotic-free medium (14). For different bacteria/antibiotic pairs, HR was shown to result mainly from genomic amplification (34), which can revert by homologous recombination between duplications. Here, we identified only four cases of amplification. The duplication of the CTX-M-15 plasmid region carrying fitness functions might have favored bacterial survival and be selected among colonies from the C-pattern (persistent-like). Two amplifications were associated with a decreased susceptibility to IMP. In the strain AC376, we observed a 89,584 bp duplication encompassing the *dcw* locus encoding major functions for cell wall synthesis and cell division. Interestingly, the duplication was not in tandem, but corresponded to an ectopic copy/paste of the region at position 688,936 (Fig. 5). The insertion was located in the *ygiW* - *qseBC* intergenic region within *ygiW* ribosome binding site (RBS). Interestingly, AC376 showed a 16-fold decrease of IMP susceptibility but almost no effect on MEM susceptibility. This observation is in agreement with previous studies showing that amplification of the *ftsQAZ* genes, part of the *dcw* locus, led to decreased susceptibility to mecillinam, an antibiotic targeting exclusively PBP2 (54). Indeed, both IMP and MEM have a high affinity to PBP2, but MEM has also some further affinity to PBP3 (40). In this case, the amplification and the phenotype were stable. Increased copy number of *bla*_KPC_ was shown to lead to a higher MIC to carbapenems (10). In strain AC150-A, we observed a 68 kb duplication within pKPC leading to a modest increase of IMP and MEM MICs (Fig. 5B, Table S4). In addition, the 11 isolates with the highest pKPC coverage belong to pattern A but, except for AC150-A, no change of IMP and MEM MICs was observed. The higher copy number might have enhanced the survival on plates supplemented with IMP but it is predicted to be unstable and likely not detectable in the experimental set-up we used.

For mutations leading to a high fitness cost, we observed frequent reversion of the growth defect and the increased MICs. A typical case was strain AC90-A from the A pattern, which showed MICs identical to the parental strain MICs and was mutated in both *cpxA* and in *cpxR*. The *cpxA* mutation would have been first selected, leading to an hyperactivation of CpxR and in turn to an overexpression of *acrB, acrD* and *eefB* efflux systems genes and a repression of the porin gene *ompK36* (32). However, the observed decreased susceptibility to carbapenems was associated with a high fitness cost with the rapid accumulation of suppressor mutations. The secondary mutation in *cpxR*, leading to an increased susceptibility compared to the parental strain, suppresses the effect of the cpxA mutation and compensates for the high cost of the primary mutation. This situation correspond to HR resulting of a mutations with a high fitness cost suppressed by a secondary mutation also reverting the effect on antibiotic susceptibility (14).

## Conclusion

HR to IMP in KPC-Kp is the result of a combination of different mechanisms to escape bactericidal antibiotics: collective protection effects, tolerance, persistence and resistance. Inter-strain variations leading to different HR frequencies across the clinical isolates here analyzed reflect the different bacterial response to IMP stress and is likely due to genetic differences that remain to be characterized. The analysis of the bacterial response to antibiotics on solid media compared to liquid culture allows to address the diversity of phenotypes and genotypes of surviving bacteria. Although the majority of colonies did not show any genetic change compared to the parental strains, we observed mutations in a very broad range of functions considerably extending our knowledge of *K. pneumoniae* intrinsic ß-lactam resistome. We hypothesize that mutations affecting these functions might also contribute to IMP treatment failure in patients infected by KPC-Kp isolates.

## Materials and METHODS

### *Klebsiella pneumoniae* isolates, media and growth conditions

The isolates analyzed in this work and their main characteristics are listed in Table S1. It includes 62 KPC-Kp clinical isolates and for comparison, 20 clinical isolates expressing other carbapenemases (OXA-48, n=3; VIM-1, n=11; NDM-1, n=3; isolates co-carrying two carbapenemases, n=3) and three isolates not carrying any carbapenemase gene as control. Bacteria were grown in Mueller-Hinton (MH), Tryptic-Soy Broth (TSB) or Lysogeny broth (LB) and on Tryptic-Soy agar (TSA) or LB agar (LBA) plates. Anaerobic growth was performed in an anaerobic chamber (Genbox BioMérieux SA, Marcy/Etoile, France) by following the manufacturer instructions. Antibiotic susceptibility testing (AST) was performed by Etest (BioMérieux) on Mueller-Hinton Agar (MHA) plates inoculated by flood with 2 ml of bacterial culture (10^6^ bacteria/ml). Flood inoculation was selected instead of swab streaking to provide a more uniform inoculation. Imipenem and meropenem breakpoints were according to the EUCAST guidelines (https://www.eucast.org/) Susceptible, ≤2 mg/L and Resistant, ≥ 4 mg/l. Etest experiments also allowed a preliminary screening for HR. We referred to HR as the occurrence of at least 5 colonies within the inhibition halo after 16-20 hours of incubation at 37 °C. Isolates with 3 or 4 colonies were defined as “moderately HR” and isolates with less than 3 colonies in the halo were classified as “non HR”. Isolates showing full contact with the Etest gradient strip for which HR could not be detected were excluded from the analysis (Table S1). Plasmid pKPC stability in strain CNR146C9 was determined by quantifying growth of 194 independent colonies isolated on LB plates in LB medium supplemented with 0.5 mg/l MEM. Strain AC420-C which have lost pKPC was used as non-growing control.

### Population Analysis Profiling (PAP)

Heteroresistance was quantified by PAP, following spreading on agar plate of either stationary phase (SP) bacterial cultures (overnight culture, ON) or exponentially growing bacteria (ON culture diluted 1000 times in TSB and incubated at 37°C until OD_600_∼0.3-0.4 was reached). Bacterial cultures were serially diluted in saline buffer and 100 μl of dilution at OD_600_ of 0.01 (an estimate of 10^7^ bacteria/ml) were spread evenly using the automatic plater EasySpiral (Interscience) on freshly prepared TSA plates containing IMP at 4-fold, 8-fold and 10-fold the MIC of each isolate, as determined by Etest. The plates were incubated at 37 °C, and colonies were counted at 24 hours., The frequency of survival was calculated as the ratio of CFUs on IMP supplemented medium over CFUs on media without IMP. Given the uncertainty in determining CFUs, we arbitrary set a threshold of 10 colonies per plate (10^2^ CFU/ml) for significance. Results were calculated from five independent experiments and expressed as the log_10_ of the frequency.

### ScanLag assay and data analysis

ScanLag assays were performed as described previously (24), but adapted to our purposes: approximately 1-3 × 10^6^ cells from mid-EP or SP cultures were spread onto TSA plates supplemented with IMP using the EasySpiral plater. Colony tracking was carried out every 20 minutes for 48-hours at 37 °C. Time of appearance (ToA) and colony growth rates were estimated by running MATLAB scripts publicly available at https://github.com/iosonofabio/scanlag (55).

Colony ToA distributions were analyzed by using the R package *mixtools* (56). We applied a two-component mixture model to the ToA distributions of both untreated populations and those surviving IMP exposure using expectation–maximization (EM) algorithm by the *normalmixEM* function. Briefly, we jointly estimated the vectors of means μ (estimated ToA mean in each component) and standard deviations Σ as well as the mixing weights λ (corresponding to the initial proportion of colonies, expressed in %) to analyze and quantitatively compare subpopulations across the selected isolates. The delay in growth of IMP exposed populations compared to untreated colonies, hereto referred to as Lag time extension (LE), was quantified as the ratio of mean ToA of N1 cells (first appearing population on IMP plates) over the mean ToA of untreated colonies. The “persister” fraction was estimated using thresholds as previously described (28). Briefly, colonies were considered as deriving from persister cells if they appeared later than three standard deviations away from the mean ToA of the N1 population (ToA > μ_N1_+3Σ_N1_). Persister frequency over the whole population was calculated per each isolate; data were then visualized using *ggplot2* (57).

### Isolation of surviving colonies

Surviving colonies were selected from IMP-containing plates, re-isolated on media containing the same IMP concentration and, in parallel, on drug free media. The next day, colonies were picked from IMP plates or from plates without IMP - if the colony did not grow on the IMP plate, and incubated in TSB without antibiotic until mid-exponential phase was reached. The cultures were then used for AST by Etest and DNA extraction for genome sequencing.

### Whole-genome sequencing and analysis

Genomic DNAs were extracted by using the DNeasy Blood & Tissue Kit (QIAGEN). *K. pneumoniae* genomes were sequenced by using the Illumina HiSeq2500 or NextSeq500 platform, with 100- and 75-nucleotides (nt) paired-end reads respectively. Libraries of *K. pneumoniae* clinical isolates were constructed by using the Nextera XT kit (Illumina) and libraries of CNR146C9 clones selected during ScanLag experiments by using NEBNext Ultra II FS DNA Library prep kit, which provide a more even genome coverage (58). Paired-end reads were assembled into contigs with SPAdes 3.9.0 (59). The complete genome sequence of CNR146C9 was determined by using the long-read PacBio technology. Accurate consensus reads obtained from raw reads with pbccs (https://github.com/nlhepler/pbccs) were assembled with Canu (60). The assembly was polished with Circulator (61) and manually corrected by mapping Illumina reads with Breseq 0.33.2 (62). Variants compared to CNR146C9 complete genome sequence were identified by using Breseq (62). Missing regions and recombination events were identified by analysing the “Unassigned missing coverage evidence” and “Unassigned new junction evidence” sections of the Breseq outputs respectively. Polymorphisms were visually verified by using Tablet (63). Genome sequences were annotated with Prokka 1.14.5 (64) and analysed for MLST and ARG content by using Kleborate (65) and Resfinder 4.0.1 (66). Plasmid incompatibility groups were identified by using PlasmidFinder 2.1 (67). Copy number variation was estimated by counting the number of reads per gene and comparing with number of Illumina reads of the parental strain CNR146C9 by using the DESSeq2 R Package (68).

### Statistical analyses

All statistical tests and graphical representations were performed using R 4.0.3 with R studio 1.3.1093 (69). All statistical tests and graphical representations were performed using R v.4.0-3 (69) using packages *ggplot2* (57) and *ggpubr* (https://rpkgs.datanovia.com/ggpubr/index.html). The log10-transformed distributions of survival frequency for exponentially-growing and stationary phases were compared using Student’s T-test. A *p-value* below the 5% threshold was deemed significant.

## Availability of data

Sequence data for the clinical isolates have been deposited at DDBJ/EMBL/GenBank (Bioproject ERP133503). CNR146C9 WGS was deposited with the following accession numbers: ERS9645073. Biosamples for the Illumina sequence data are listed in Table S1.

## ACKNOWLEDGEMENTS

This work was supported by grants from the French National Research Agency IBEID (ANR-10-LABX-62-IBEID) and SEQ2DIAG (ANR-20-PAMR-0010). Adriana Chiarelli is part of the Pasteur - Paris University (PPU) International PhD Program, with funding from the Institut Carnot Pasteur Microbes & Santé, and the European Union’s Horizon 2020 research and innovation programme under the Marie Sklodowska-Curie grant agreement No 665807.

The authors thank Virgile Andreani and Gregory Bath for discussion on HR, Guilhem Royer for critical reading of the manuscript, Laurence Ma from the Institut Pasteur Biomics platform (C2RT, Institut Pasteur, Paris, France, supported by France Génomique, ANR-10-INBS-09-09 and IBISA) for her help with Illumina sequencing.

---

**Fig. S1.**
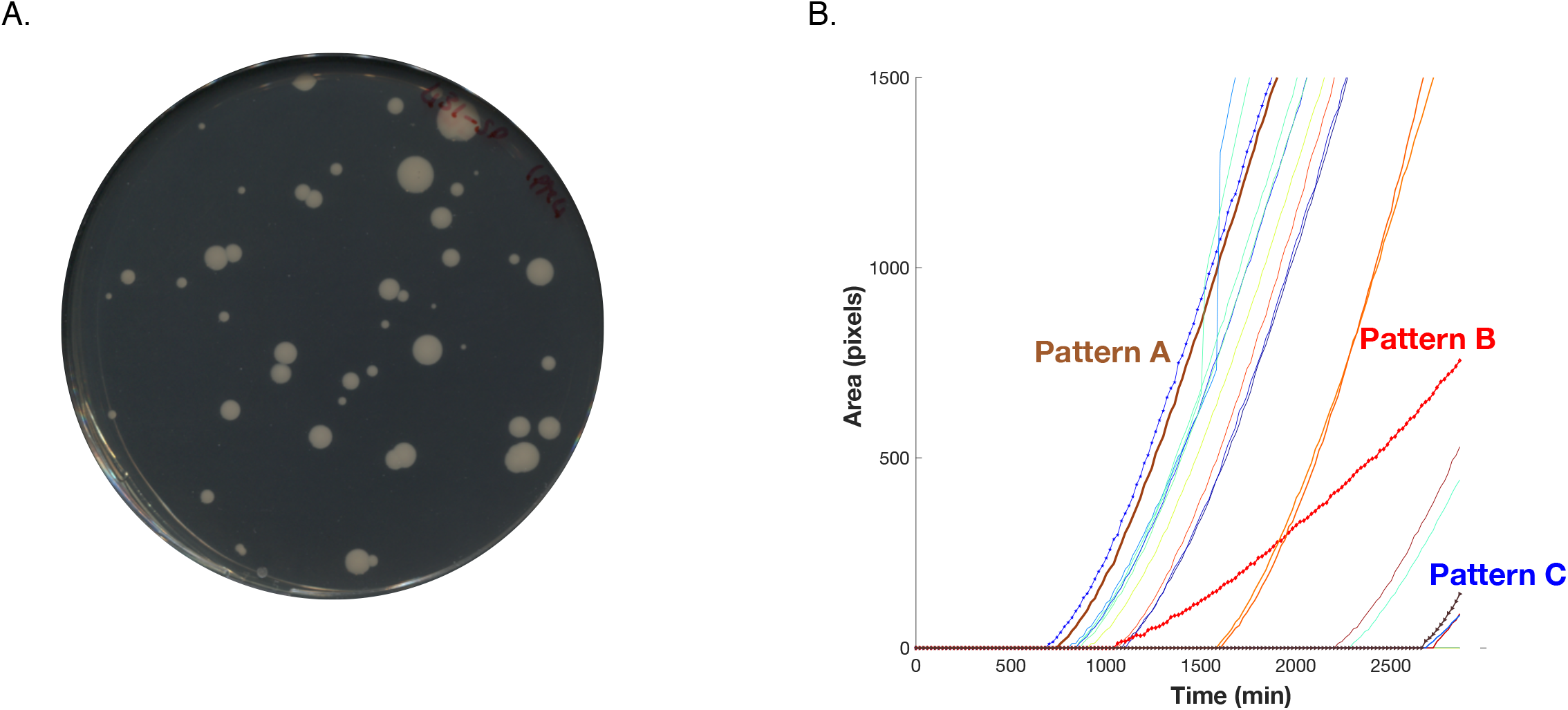
Heterogeneity of colony growth as detected by ScanLag. **(A)** Example of plate containing colonies from the isolate CNR146C9 emerging on media supplemented with imipenem at 8-fold the MIC as determined by Etest. Colonies exhibit heterogeneous size and morphology, reflected by different Time of Appearance and Growth rates. **(B)** Colony growth curves as determined by ScanLag. Three patterns (A, B and C) were described according to the time of appearance and the growth rate.

**Fig. S2.**
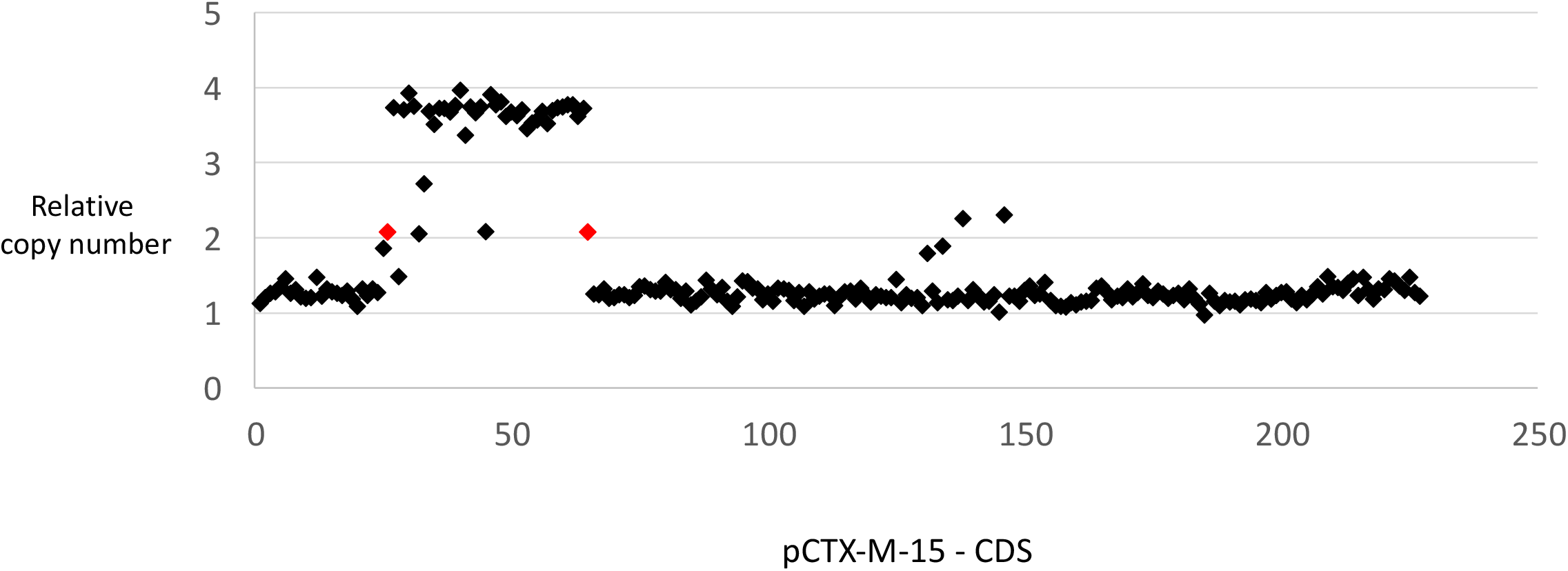
DNA amplification in pCTX-M-15. Duplication of a c.a. 49 kb region within the pCTX-M-15 plasmid in strain AC362-C. Average copy number of the 227 CDS of pCTX-M-15 was calculated relative to the corresponding average value for all isolates. Genes are indicated by diamonds; in red, the two IS*5075* bracketing the duplication.

