## Supplemental Table 3 for "Carbapenem heteroresistance of KPC-producing *Klebsiella pneumoniae* results from tolerance, persistence and resistance"

**Table S3. CNR146C9 chromosome, plasmids and antibiotic resistance genes**

| | | Chromosome | pCTX-M-15 | pKPC | Circular phage |
|---|---|---|---|---|---|
| replicon | | | IncFIB/IncFII | IncFIB(pQil)/IncFII | Phage |
| size (base pairs) | | 5351626 | 237007 | 110489 | 52608 |
| ARG | Aminoglycoside | | aac(6')-Ib-cr<br>aac(3)-IIa<br>aph(3'')-Ib<br>aph(6)-Id | | |
| | ß-lactam | $bla_{SHV-28}$* | $bla_{TEM-1B}$<br>$bla_{OXA-1}$<br>$bla_{CTX-M-15}$ | $bla_{TEM-1A}$<br>$bla_{KPC-3}$<br>$bla_{OXA-9}$ | |
| | Fluoroquinolone | *oqxA*<br>*oqxB* | *qnrB1* | | |
| | Fosfomycin | *fosA* | | | |
| | Phenicol | | *catB2* | | |
| | Sulfonamide | | *sul2* | | |
| | Trimethoprim | | *dfrA14* | | |

* SHV-28 like;
